# Transcriptomic profile of *MTUS1* -low TNBC reveals candidate therapeutic strategies

**DOI:** 10.64898/2026.05.22.727134

**Authors:** Gwenn Guichaoua, Olivier Collier, Sylvie Rodrigues-Ferreira, Clara Nahmias, Véronique Stoven

## Abstract

**Background:** Triple-negative breast cancer (TNBC) is a clinically aggressive breast cancer subtype. It is a heterogeneous disease that remains difficult to stratify and that still lacks durable and biomarker-guided therapeutic options. Low expression of the tumour suppressor *MTUS1* is associated with aggressive breast cancer features, but the biological properties of *MTUS1*-low TNBC remain insufficiently defined. Our goal was to determine whether low *MTUS1* expression defines shared proliferative and stress-adaptation mechanisms that could guide candidate therapeutic strategies and corresponding target/drug pairs in *MTUS1*-low TNBC.

**Methods:** We labelled tumours from seven public TNBC RNA-seq cohorts based on the lowest and highest *MTUS1* expression tertiles. Differential gene expression was analysed using gene set enrichment analysis (GSEA) on the Hallmark pathway database to identify deregulated biological pathways between *MTUS1*-low TNBC tumours and their *MTUS1*-high counterparts. Reproducibility was examined across independent TNBC cohorts and secondarily in broader breast cancer and selected TCGA tumour cohorts. Gene essentiality scores from CRISPR–Cas9 experiments in TNBC cell-line models were correlated to *MTUS1* expression in these cell lines, to propose therapeutic strategies and their corresponding candidate target/drug pairs.

**Results:** *MTUS1*-low tumours showed a reproducible pathway-level proliferation mechanism driven by the MYC oncogene and sustained by up-regulated oxidative phosphorylation, combined with stress adaptation mechanisms involving unfolded protein response (UPR), and DNA repair Hallmark gene sets. Based on CRISPR data, we propose 3 therapeutic strategies: (1) targeting MYC to reduce its transcriptional activity, (2) targeting proteins from UPR, (3) targeting DNA-repair. We also propose corresponding candidate target/drug pairs to allow experimental validation of these strategies.

**Conclusions:** Proliferation in low *MTUS1* TNBC is driven by MYC and stress-adaptation mechanisms. By linking this tumour profile to CRISPR-derived dependency signals, our analysis prioritises experimentally testable target–pathway hypotheses centred on MYC, UPR/proteostasis, and DNA-repair or checkpoint control. Although the proposed therapeutic strategies and candidate targets remain to be experimentally tested, the latter finding is consistent with published work showing that ATIP3-deficient TNBC cell line models are sensitive to inhibition of the WEE1 PKMYT1 G2/M checkpoint kinases.

## 1 Introduction

Triple-negative breast cancer (TNBC), defined by the absence of oestrogen receptor (ER), progesterone receptor (PR), and HER2 expression, is a clinically aggressive breast cancer subtype. Immune checkpoint blockade in selected PD-L1-positive tumours and PARP inhibitors in germline *BRCA1/2*-mutant tumours have improved outcomes for some patients [1], but many TNBCs still lack durable and biomarker-guided therapeutic options. This unmet need reflects, in part, the molecular heterogeneity of TNBC and motivates the identification of specific tumour profiles that capture underlying aggressive biology and guide mechanistic follow-up [2].

Large-scale transcriptomic and functional genomic resources now make it possible to connect tumour profiles observed in patients with candidate targets measured in experimental models [3–5]. Such integration is useful when a biomarker is associated with a poor outcome.

In this context, the *MTUS1* locus (8p22) has been proposed as a tumour suppressor region in breast cancer and other malignancies [6]. Reduced *MTUS1* expression has been reported across multiple cancer types, including breast cancer, and has been associated with adverse clinicopathological features and outcomes [7–9]. *MTUS1* encodes several splice isoforms, including ATIP1, ATIP3, and ATIP4, but in breast cancer, ATIP3 has been characterised as the main isoform [10]. ATIP3 is a microtubule-associated regulator of mitotic spindle organisation and chromosome segregation [7, 10, 11], supporting a role in mitotic control and tumour aggressiveness. It is important to note that downregulation of *MTUS1* /ATIP3 has been reported in the majority of TNBC tumours [7, 10]. However, prior knowledge has not yet established broader biological characteristics or transcriptional programmes that would be specific to *MTUS1*-low TNBC.

In the present paper, based on transcriptional data from seven independent TNBC cohorts, we identify deregulated biological pathways that characterise TNBC tumours with low *MTUS1* expression. Based on these pathways, we use CRISPR data from TNBC cell-line models to propose three therapeutic strategies and suggest corresponding candidate target/drug pairs that will allow further experimental validation.

## 2 Materials and Methods

### 2.1 Transcriptomic characterisation of *MTUS1*-low TNBC

#### 2.1.1 Patient cohorts

##### TNBC cohort datasets

We considered seven independent cohorts of primary TNBC tumours for which gene-level raw read counts were available, as shown in Table 1. A pooled dataset was also generated for integrative analyses, as described below. VUMC raw counts were provided by the authors [12]. For GEO cohorts (GSE181466, GSE192341, and GSE202203), gene-level raw count matrices and clinical annotations were retrieved from GEO using the `GEOquery` package [13]. TCGA_BRCA raw counts and clinical annotations were retrieved from the NCI Genomic Data Commons (GDC) via its API (https://api.gdc.cancer.gov/) [14]. For SRA-based cohorts (SRP042620 and SRP157974), sample annotations and gene-level coverage counts were obtained from the `recount3` resource. Counts were converted to gene-level integer count matrices using the `transform_counts()` function, following `recount3` recommendations for differential expression analyses [15]. The VUMC dataset [12], GSE181466, and SRP157974 comprised TNBC samples exclusively. The remaining cohorts (GSE192341, GSE202203, SRP042620, and TCGA_BRCA) include multiple breast cancer (BC) subtypes, and molecular subtype annotations were used to extract TNBC.

**Table 1:** RNA-seq datasets used in this study, providing the number of breast cancer (BC) and triple-negative breast cancer (TNBC) samples in each cohort, including the pooled dataset. Asterisks (*) denote cohorts composed exclusively of TNBC samples.

| Dataset | Number of BC samples | Number of TNBC samples |
| --- | --- | --- |
| SRP042620 | 84 | 42 |
| VUMC* | 44 | 44 |
| GSE192341 | 87 | 52 |
| GSE181466* | 79 | 79 |
| TCGA_BRCA | 1 111 | 116 |
| GSE202203 | 2 913 | 291 |
| SRP157974* | 448 | 448 |
| All_pooled | 4 766 | 1072 |

**Table 2:**
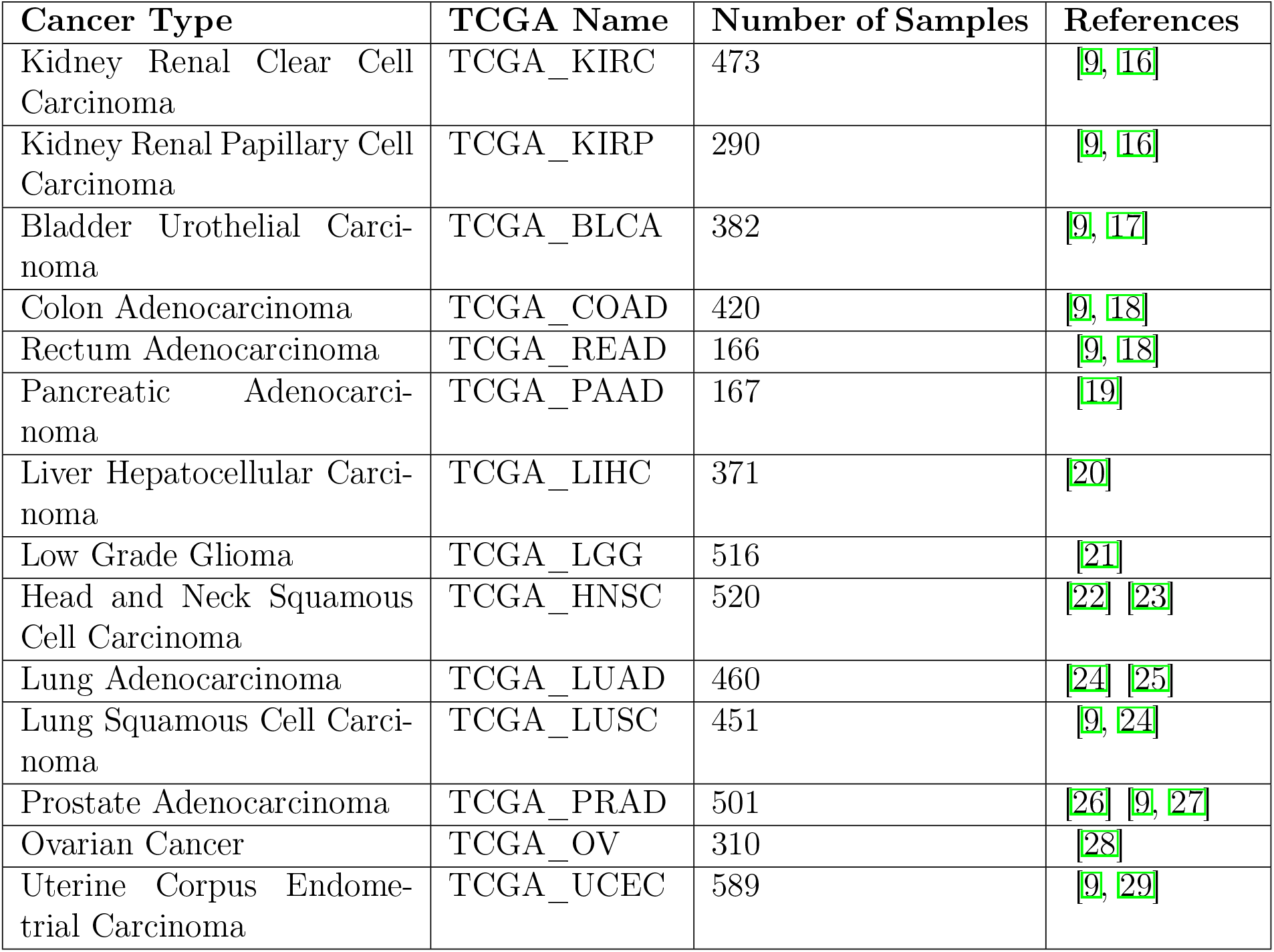
RNA-seq datasets of multiple cancer locations from TCGA cohorts used in this study.

##### BC cohort datasets

Because three of the seven TNBC cohorts included fewer than 52 samples, analyses restricted to these datasets could be more sensitive to tumour heterogeneity or patient-specific variability than to reproducible effects associated with low *MTUS1* expression. To mitigate this limitation and increase statistical power, we additionally analysed the corresponding full breast cancer (BC) cohorts from which TNBC cases were extracted, when applicable.

##### Selected non-breast TCGA cohorts

We also analysed selected tumour types from The Cancer Genome Atlas (TCGA). Cohorts were selected based on prior reports indicating (i) an association between low *MTUS1* expression and poorer patient outcome and/or (ii) reduced *MTUS1* expression in tumour versus matched normal tissues [9]. TCGA RNA-seq gene-level raw read counts and associated clinical annotations were retrieved from the NCI Genomic Data Commons (GDC) via its API (https://api.gdc.cancer.gov/). Table summarises the selected cancer types, corresponding TCGA projects, sample sizes, and supporting references.

#### 2.1.3 Differential expression and pathway enrichment

Differential analyses were conducted to quantify gene-level expression changes and to identify biological pathways consistently associated with low *MTUS1* expression (*MTUS1* - low) relative to high *MTUS1* expression (*MTUS1*-high) tumours.

##### Labeling tumours based on *MTUS1* expression

For each RNA-seq dataset, let *C* = (*C*_*j,k*_) ∈ ℕ ^*J*×*K*^ denote the gene-level count matrix, where *J* is the number of genes and *K* the number of samples. Lowly expressed genes were filtered within each dataset by removing genes with total counts across all samples ≤ 10. To define *MTUS1*-low and *MTUS1*-high groups, samples were stratified using within-cohort tertiles of DESeq2-normalised *MTUS1* expression. Samples in the lower and upper tertiles were assigned to the *MTUS1*-low and *MTUS1*-high groups, respectively, whereas samples in the intermediate tertile were excluded from downstream differential analyses, to increase contrast between groups and reduce ambiguity around borderline expression values. This method was used to define *MTUS1*-low and *MTUS1*-high groups in all cohorts in the present study (TNBC, BC and TCGA cohorts).

##### Pooled datasets

Pooled datasets were built by concatenating samples across cohorts, keeping only genes shared by all datasets. Sample-level metadata were merged by column alignment. *MTUS1*-low/*MTUS1*-high labels were defined within each cohort before pooling and subsequently carried into pooled analyses, to limit cohort-driven thresholding. For pooled analyses, the cohort of origin was included as a covariate in the design formula to account for between-study effects. In addition, only consortium- or study-provided technical batch variables were considered as adjustment covariates (e.g. TCGA internal batch annotations, SRP157974 batch fields when available); no surrogate or data-driven batch factors were inferred.

##### Gene-level differential expression

Gene-level differential expression was computed using `PyDESeq2` [30]. For each gene *j*, `PyDESeq2` estimated the log_2_ fold change LFC_*j*_ for *MTUS1*-low versus *MTUS1*-high, together with the Wald statistic *W*_*j*_ and the corresponding Benjamini–Hochberg adjusted *p*-value *p*_*j*_ [31]. When applicable, batch variables were included as covariates in the model design. Gene-wise Wald statistics were used as input for pathway-level analyses.

##### Hallmark pathway enrichment

Pathway enrichment was assessed using a pre-ranked gene set enrichment analysis (GSEA) implemented in `decoupleR` [32]. Genes were ranked by Wald statistics *W*_*j*_; for each gene set *P*, enrichment was summarised by a normalised enrichment score NES(*P*) and an FDR-adjusted *p*-value *p*_*P*_. Analyses focused on the 50 MSigDB Hallmark gene sets [33, 34].

##### Reproducibility criteria

Hallmark pathways were selected using prespecified reproducibility criteria applied on the seven TNBC cohorts: (i) consistent NES signs across all cohorts, and (ii) FDR < 0.05 in at least six of the seven cohorts, allowing up to one cohort with FDR ≥ 0.05.

### 2.2 Identification of *MTUS1*-associated essential genes in TNBC cell lines

#### 2.2.1 Gene functional dependency data

The DepMap Public v24Q4 release [35], accessed through the DepMap portal [36] and developed within the Cancer Dependency Map programme [37], provides a large collection of human cancer cell lines, including transcriptomic data and functional dependency scores derived from genome-wide CRISPR-Cas9 loss-of-function screens. We selected the 30 cell lines annotated as TNBC. Gene expression data were obtained from DepMap RNA-seq profiles provided as log_2_(TPM+1) matrices, and *MTUS1* expression values were taken from DepMap RNA-seq profiles to ensure a consistent expression scale across resources. Gene dependency data were retrieved from the same release and quantified using the Chronos algorithm [5], which models population dynamics following gene knockout. Chronos provides a gene-level essentiality score for each cell line, where values near 0 denote non-essential genes and common essential genes typically show median scores around −1. From the 30 TNBC cell lines, we retained those for which RNA-seq and Chronos profiles were both available (*n* = 24; cell-line identifiers are reported in Supplementary Table S2).

#### 2.2.2 Correlation between *MTUS1* expression and gene essentiality

Given the limited number of TNBC cell-line models and the use of continuous *MTUS1* expression values in DepMap (log_2_(TPM + 1)), we adopted a correlation-based strategy to relate gene essentiality directly to *MTUS1* expression across TNBC cell lines rather than stratifying models into *MTUS1*-low/*MTUS1*-high tertiles. Among 18,444 genes, we first restricted the analysis to genes showing evidence of essentiality in TNBC (Chronos score < *-*1 in at least one TNBC cell line; *n* = 1,854) to reduce noise from uniformly non-essential genes. For each of these genes, we computed the Pearson correlation between Chronos gene effect scores and *MTUS1* expression across the matched TNBC models. Because more negative Chronos scores indicate stronger dependency, a positive correlation indicates increased dependency at lower *MTUS1* expression. Spearman correlations were computed as a robustness check and are reported in Supplementary Table S3. Candidate essential genes were prioritised using nominal correlation *p*-values (*p* < 0.05), corresponding approximately to |r| ≳ 0.40 in two-sided tests. The full distribution of correlation coefficients and the relationship between correlation magnitude and significance are shown in Figure 3A–B. Given the limited number of TNBC models and the resulting lack of power for genome-wide multiple-testing correction, this analysis was treated as exploratory and hypothesis-generating.

#### 2.2.3 Functional analysis of *MTUS1*-associated essential genes according to up-regulated Hallmark pathways in *MTUS1*-low TNBC

Genes showing nominally significant positive correlations (increased dependency with decreasing *MTUS1* expression) that mapped to the deregulated Hallmark pathways highlighted in the transcriptomic study performed on *MTUS1*-low TNBC ((Section 2.1.2)) were prioritised as candidate targets in these pathways. To quantify enrichment of pathways among *MTUS1*-associated dependencies, we performed over-representation analysis (ORA) using the hypergeometric test implemented in `gseapy (gp.enrichr)` [38]. The background universe was defined as the set of genes tested in the correlation analysis (*n* = 1,854). Enrichment *p*-values were adjusted for multiple testing using the Benjamini–Hochberg procedure, and pathways with FDR-adjusted *p* < 0.05 were considered significant.

## 3 Results

We first defined the transcriptional modifications and functional pathways associated with *MTUS1*-low TNBC tumours. Then, based on CRISPR dependency data, we examined whether these pathways contain genes whose functions are critical for survival in TNBC cell-line models.

### 3.1 *MTUS1*-low TNBC are characterized by specific transcriptional programs

Tumours were first stratified as *MTUS1*-low or *MTUS1*-high using within-cohort *MTUS1* expression tertiles, as described in Section 2.1.2. Given cohort heterogeneity and the limited sample size of some datasets, we prioritised pathway-level rather than gene-level analysis to capture coordinated biological shifts and improve cross-study interpretability [39]. We chose the 50 MSigDB Hallmark gene sets [33] as a reference gene set database, because Hallmark summarises major cancer-relevant cellular processes. For each TNBC cohort, GSEA compared *MTUS1*-low with *MTUS1*-high tumours. Twelve Hallmark pathways met the prespecified reproducibility criteria: concordant NES signs across all cohorts and FDR < 0.05 in at least six of seven cohorts (Figure 1A). The pooled TNBC cohort showed concordant results but was used only as secondary support for the analyses, not for pathway discovery.

**Figure 1:**
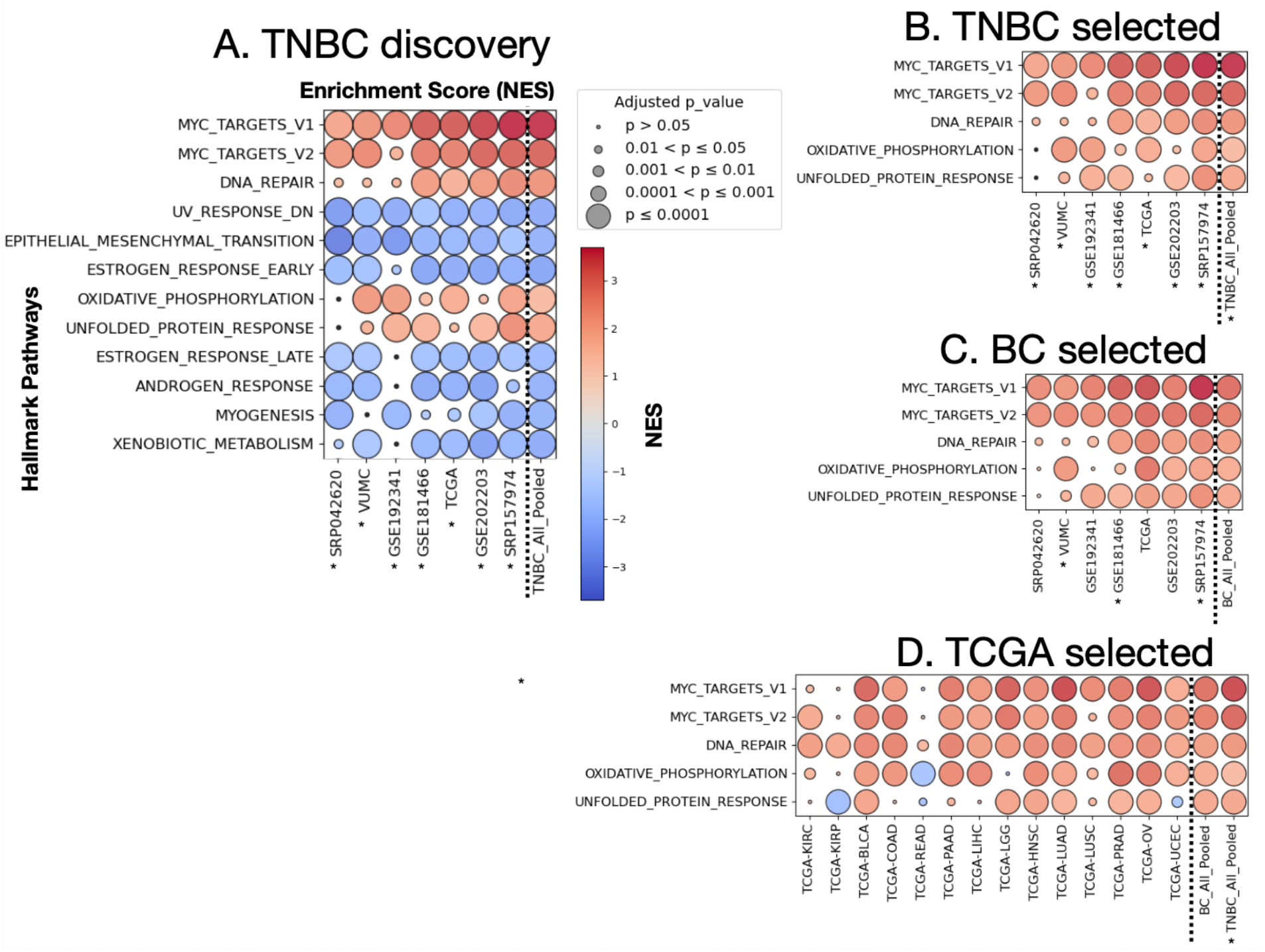
Hallmark pathway enrichment in *MTUS1*-low versus *MTUS1*-high tumour cohorts. Circles are coloured by normalised enrichment score (NES; red: enrichment in *MTUS1*-low, blue: depletion in *MTUS1*-low) and scaled by the FDR-adjusted pathway *p*-value. Asterisks (*) denote cohorts composed exclusively of TNBC samples. Dotted separators indicate pooled datasets. (A) Discovery of Hallmark pathways meeting the reproducibility criteria in seven TNBC cohorts (concordant NES signs in all cohorts, FDR < 0.05 in at least six out of seven cohorts). (B) Reproducible pathways enriched in *MTUS1*-low TNBC tumours (NES> 0). (C) Evaluation of the selected pathways in broader BC cohorts comprising all BC subtypes. (D) Evaluation of the selected pathways in TCGA tumour cohorts from other anatomical sites and in pooled cohorts.

Among the twelve transcriptionally deregulated Hallmark pathways in *MTUS1*-low tumours (NES> 0), five were consistently up-regulated and were kept for further analysis (Figure 1B). Indeed, our aim was not only to characterise *MTUS1*-low TNBC tumours, but also to identify candidate therapeutic strategies and targets. In this context, inhibition of an overexpressed/overactivated protein target is a more classical and achievable approach than upregulation of the expression or of the function of a down-regulated protein target. In addition, up-regulated pathways provided the most direct basis for comparison with gene dependencies from the CRISPR data. Notably, the five TNBC-selected pathways remained up-regulated in broader breast cancer cohorts comprising all molecular subtypes (Figure 1C), and in most additional TCGA cohorts from other tumour localisations (Figure 1D).

Thus, the *MTUS1*-low transcriptional profile in TNBC is characterised by a reproducible signature in Hallmark. MYC-linked pathways (MYC_TARGETS_V1/V2) dominated the signal, together with `UNFOLDED_PROTEIN_RESPONSE`, `DNA_REPAIR`, and `OXIDATIVE_ PHOSPHORYLATION` pathways. Sign preservation outside the TNBC discovery cohorts supports the robustness of this profile.

### 3.2 Proliferation in *MTUS1*-low TNBC is driven by MYC and supported by energy and stress adaptation pathways

#### Up-regulation of MYC-related pathways

In *MTUS1*-low TNBC, the strongest and consistent signals were observed for the two MYC-associated Hallmark gene sets, although *MYC* mRNA itself was only modestly increased in *MTUS1*-low tumours (0 < log_2_ fold change < 1 across datasets), with statistical significance mainly in the largest cohorts and pooled analyses (Table S1). *MYC* is a well-established oncogene and transcriptional regulator involved in growth and cell-cycle processes [40, 41]. These two gene sets capture complementary roles of MYC’s activity: `MYC_TARGETS_V1` (200 genes) reflects broad biosynthetic and proliferation-associated processes, whereas `MYC_TARGETS_V2` (58 genes) comprises oncogenic processes including cell-cycle control and tumour progression [33, 42]. Their limited overlap (18 shared genes) supports that they represent non-redundant facets of MYC-driven transcriptional activity [33]. In TCGA cohorts with other localisations, upregulation of *MYC* pathways was also globally observed, although with variable intensities, indicating a tissue-context dependence for this trait.

#### Energy production and stress adaptation pathways

A few additional pathways support MYC-driven proliferation in *MTUS1*-low TNBC. First, the `OXIDATIVE_PHOSPHORYLATION` Hallmark pathway (200 genes) is upregulated, which means increased mitochondrial activity for ATP production. While many cancers rely on aerobic glycolysis (Warburg effect) [43], oxidative phosphorylation remains critical in aggressive and therapy-resistant states [44]. This aligns with the energetic demands of MYC-driven transcriptional program and with reports showing that MYC can promote mitochondrial metabolism, including in TNBC [45–47].

Second, the `DNA_REPAIR` Hallmark pathway (150 genes) is up-regulated, suggesting enhanced engagement of genome maintenance programs to allow survival under replication stress and DNA damage in high proliferating *MTUS1*-low TNBC. In addition, increased mitochondrial respiration can elevate reactive oxygen species (ROS) [48], providing an endogenous source of oxidative DNA damage, which strengthens the DNA damage response. Consistent with this interpretation, *MTUS1* depletion has been linked to replication stress and DNA damage pathway engagement [49].

Third, `UNFOLDED_PROTEIN_RESPONSE` Hallmark pathway (113 genes) is up-regulated, indicating activation of ER stress pathways to preserve proteostasis and promote survival under proteotoxic load [50–52]. Increased protein synthesis downstream of MYC activation can challenge cells’ protein folding capacity [45], providing a rationale for UPR activation in *MTUS1*-low tumours.

### Upregulated pathways in *MTUS1*-low TNBC cooperate in a functional network

As presented above, high proliferation rates in *MTUS1*-low TNBC appear to be driven by MYC-associated pathways, while the resulting increases in energy demand and replication stress are covered respectively by upregulation of the oxidative phosphorylation pathway and by upregulation of DNA-repair and UPR pathways. Therefore, the five complementary up-regulated pathways highlighted in this study can be integrated within a functional network that illustrates how these pathways cooperate, as presented in Figure 2. Overall, this model recapitulates the main functional characteristics shared by *MTUS1*-low TNBC tumours and explains the agressivity of the corresponding disease [10, 49].

**Figure 2:**
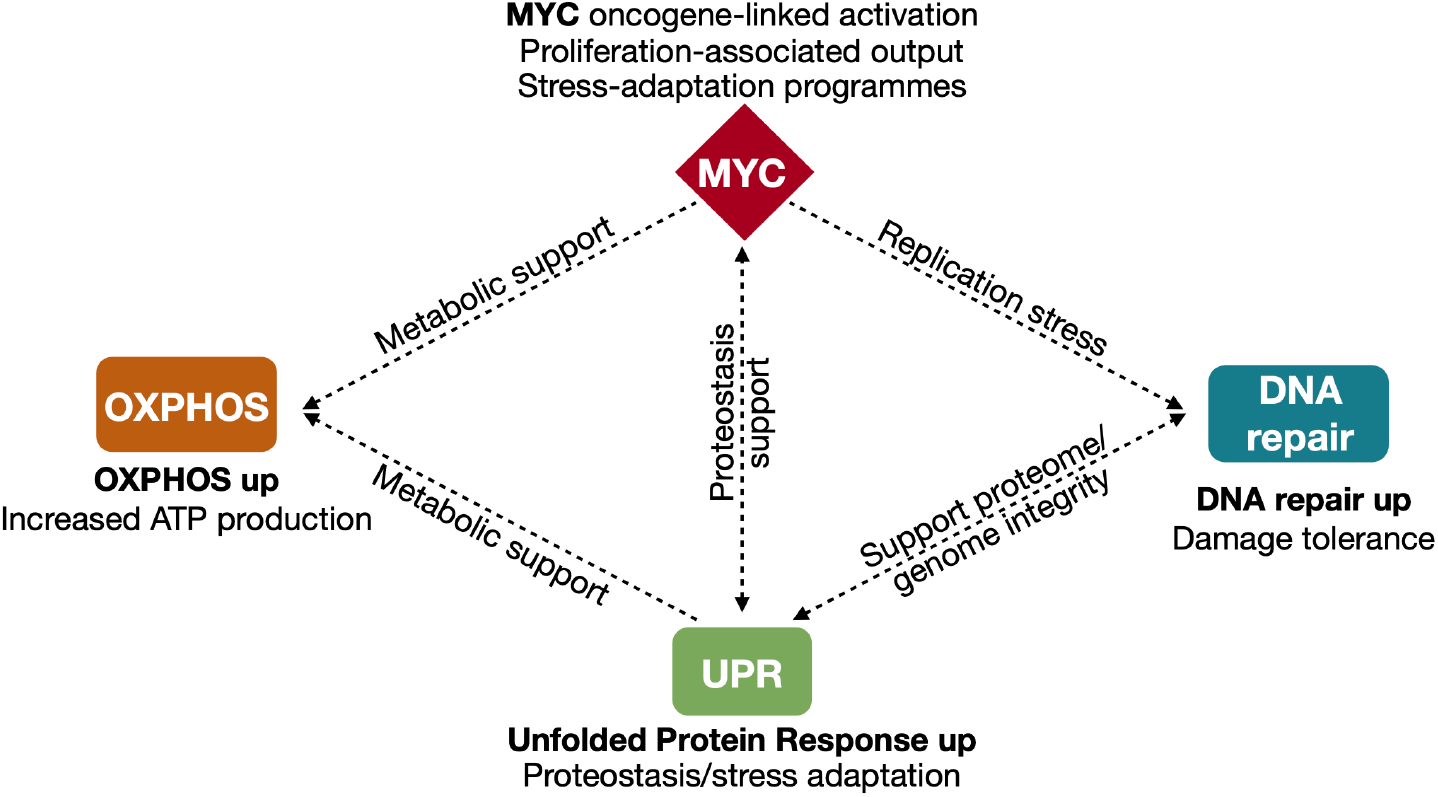
Proposed functional network integrating up-regulated Hallmark pathways in *MTUS1*-low TNBC. Dashed arrows indicate functional associations supported by pathway enrichment and published evidence rather than causal directionality inferred from the present datasets.

This network provides a tumour-derived representation of the *MTUS1*-low TNBC up-regulated biological characteristics. However, by itself, it does not establish which genes within these pathways are most critical for tumour-cell survival or could serve as candidate therapeutic targets.

### 3.3 CRISPR dependency data link genes from the *MTUS1*-low TNBC network to candidate targets

We used CRISPR dependency data performed on TNBC cell-line models, and searched for genes whose essentiality (also called dependency) increases as *MTUS1* expression decreases in these cell lines. More precisely, we analysed genome-wide DepMap v24Q4 Chronos gene effect scores (see Section 2.2.1), and because more negative Chronos scores indicate stronger essentiality, a positive correlation between Chronos score and *MTUS1* expression indicates that dependency increases as *MTUS1* expression decreases.

#### Essential genes associated with low *MTUS1*

Among 18 444 genes, we first selected those showing evidence of essentiality (Chronos score < −1) in at least one TNBC cell line, leading to *n* = 1 854 genes. The distribution of the correlations between the Chronos scores of these genes with *MTUS1* expression was centred near zero, as shown in Figure 3A). However, 87 genes fell in the positive tail of the distribution, showing increased essentiality as *MTUS1* expression decreased at nominal significance thresholds (*p* < 0.05; Figure 3B). These 87 genes were retained as *MTUS1*-associated candidate essential genes in TNBC.

**Figure 3:**
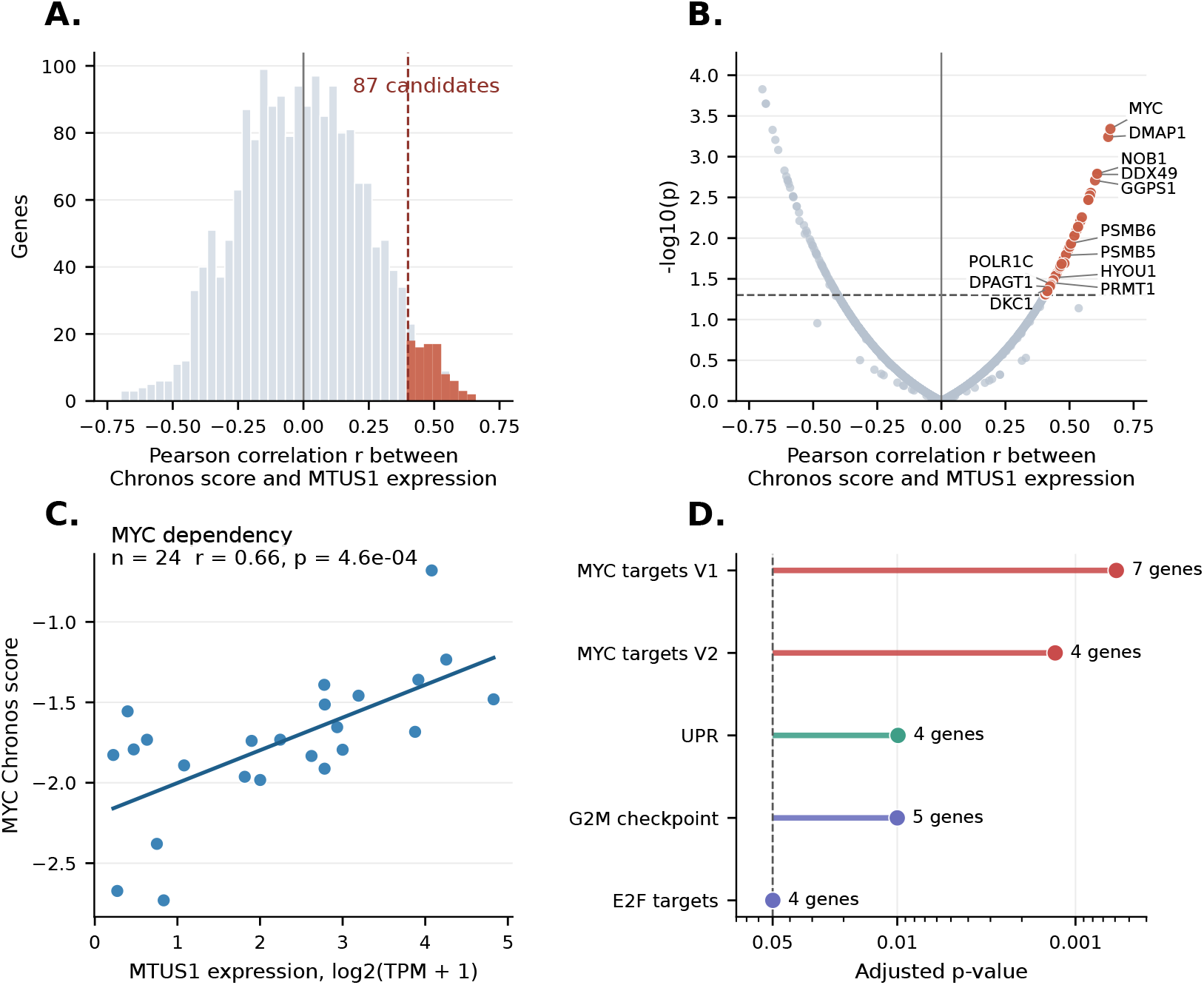
DepMap CRISPR dependency analysis. (A) Distribution of Pearson correlation coefficients between *MTUS1* expression and Chronos gene scores (i.e. essentiality score) across *n* = 1 854 essentiality-filtered genes. Positive correlations indicate stronger dependency as *MTUS1* expression decreases. The dashed red line marks the approximate nominal significance threshold corresponding to *p* < 0.05. (B) Relationship between Pearson correlation coefficients and nominal significance. Orange points indicate the 87 *MTUS1*-associated candidate essential genes. (C) Example of *MYC* as essential gene: the positive correlation between *MTUS1* expression and MYC dependency (Chronos score) indicates stronger essentiality of MYC in TNBC cell lines with lower *MTUS1* expression. (D) Hallmark pathway over-representation among the 87 candidate essential genes, using *n* = 1 854 essentiality-filtered genes as the background universe. The dashed line indicates adjusted *p* = 0.05.

Spearman correlations were computed as a robustness check and are reported in Supplementary Table S3. Notably, *MYC* was among the strongest candidates (*r* = 0.66, *p* = 4.6 × 10^−4^; Figure 3C), consistent with MYC as a key driver gene in *MTUS1*-low TNBC tumours.

#### Hallmark pathways enriched in *MTUS1*-associated essential genes

We next tested whether the 87 *MTUS1*-associated essential genes were enriched in Hallmark pathways, using the 1 854 essentiality-filtered genes as the background universe (Section 2.2.2). As shown in Figure 3D) and Table 3, we found that 5 pathways were significantly enriched, including 3 pathways belonging to the *MTUS1*-low TNBC tumours network in Figure 2: `MYC_TARGETS_V1`,`MYC_TARGETS_V2`, and `UNFOLDED_PROTEIN_RESPONSE`. This indicates that the survival of *MTUS1*-low TNBC tumours may depend particularly on these pathways. Moreover, although the `DNA_repair` pathway was not enriched in *MTUS1*-associated essential genes, it contained two *MTUS1*-associated essential genes. On the contrary, no *MTUS1*-associated essential gene was found in the `OXIDATIVE_PHOSPHORYLATION` pathway, which indicates that energy supply may not be the limiting biological process to sustain proliferation of *MTUS1*-low TNBC tumours. Overall, 16 *MTUS1*-associated essential genes in TNBC cell lines belonged to four out of five Hallmark pathways consistently up-regulated in *MTUS1*-low tumours and shown in Figure 2.

**Table 3:**
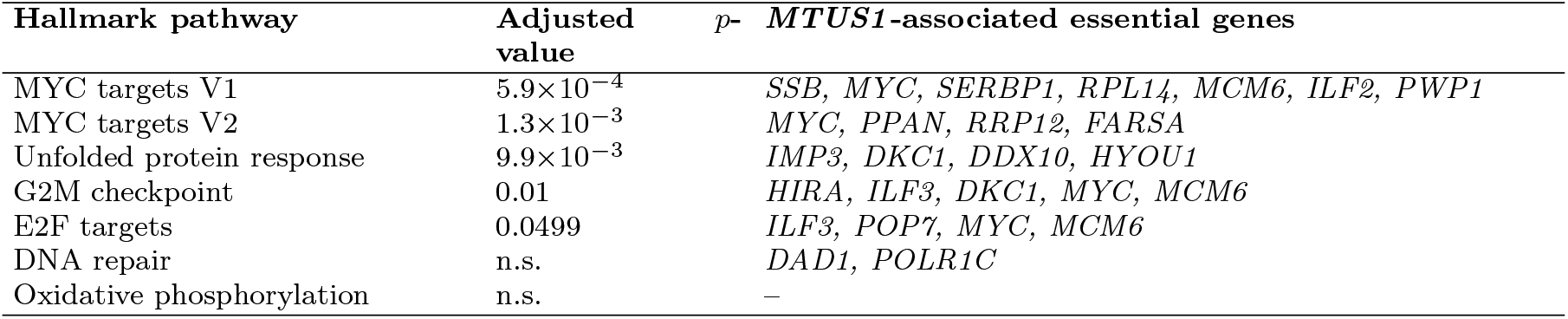
Hallmark pathways enriched in *MTUS1*-associated essential genes in TNBC cell lines. Adjusted *p*-values are from over-representation analysis using the 1 854 essentiality-filtered genes as background and Benjamini–Hochberg correction. n.s., not significant in this over-representation test.

As shown in Table 3, the `G2M_CHECKPOINT` and `E2F_TARGETS` Hallmark pathways were also enriched in *MTUS1*-associated essential genes in TNBC cell-line models. These pathways were not part of the five up-regulated pathways in *MTUS1*-low TNBC tumours, but they are functionally related to them: G2M checkpoint activity links cell-cycle progression to DNA damage and control of replication-stress [53, 54], whereas E2F target gene sets support G1/S transition, DNA replication, and DNA repair-associated transcriptional responses [55, 56].

Taken together, although TNBC cell lines are imperfect models of TNBC tumours, *MTUS1*-low up-regulated biological pathways and *MTUS1*-associated essential genes in TNBC models appear functionally consistent.

### 3.4 Candidate therapeutic targets for *MTUS1*-low TNBC

We propose to search for candidate targets among the 16 *MTUS1*-associated essential genes in Table 3 that also belong to up-regulated pathways in *MTUS1*-low TNBC tumours, i.e. to the Hallmark MYC-related, `DNA_repair`,and`UNFOLDED_PROTEIN_RESPONSE` pathways. Among these 16 genes, we kept those coding for ‘druggable proteins’, i.e. for which known pharmacological inhibitors were available (including, when possible, known marketed drugs), because this allows future experimental evaluation, for example, based on survival assays. As shown in Table 4, this left us with four main Hallmark-derived candidate targets: MYC, POLR1C, and DKC1 and HYOU1, respectively, for the MYC-related, `DNA_repair`, and `UNFOLDED_PROTEIN_RESPONSE` pathways. More generally, in terms of biological mechanisms, these candidate targets lead us to propose three experimentally testable candidate therapeutic strategies for *MTUS1*-low TNBC tumours: (1) targeting MYC to down-regulate its transcriptional program, (2) targeting proteins in the `DNA_repair` pathway to increase replication stress to untolerable levels, and (3) targeting proteins of the `UNFOLDED_PROTEIN_RESPONSE` pathway to increase ER stress to untolerable levels, and thus promoting cell death. These three axes are detailed below.

**Table 4:** Proposed therapeutic strategies and their corresponding target/drug pairs. The upper part of the Table reports candidate targets present in the Hallmark database. The lower part of the Table reports candidate targets that are not in Hallmark.

| Hallmark Pathway | Therapeutic strategy | Target | Drugs/Inhibitors |
| --- | --- | --- | --- |
| MYC_TARGETS_V1/V2 | Down-regulation of MYC activity | *MYC* | OMO-103 [57]; OTX-2002 [58]; MYCi-361, MYCi-975, and MYCMI-6 [59] |
| DNA_REPAIR | Increase replication stress | *POLR1C* | CX-5461 (RNA polymerase I inhibition) [60]; ML-60218 (RNA polymerase III inhibition) [61] |
| UNFOLDED_PROTEIN_RESPONSE | Increase ER stress | *DKC1*<br>*HYOU1* | Pyrazofurin [62]<br>HYOU1-IN-1 (Compound 33) [63] |
| Biological Process | Therapeutic strategy | Target | Drugs/Inhibitors |
| Related to DNA_REPAIR | Increase replication stress | *PRMT1* | CTS2190 [64] |
| Related to UPR | Increase ER stress | *DPAGT1*<br>*PSMB5*<br>*PSMB6* | Tunicamycin V; muraymycin A1 [65]<br>Bortezomib; carfilzomib<br>Bortezomib; carfilzomib |

#### Targeting MYC to down-regulate its transcriptional program

The strongest signal in *MTUS1*-low TNBC involved MYC_TARGETS_V1/V2, while *MYC* itself emerged among the strongest *MTUS1*-associated essential genes, even when *MYC* mRNA up-regulation was modest. These results support the interest of testing whether *MTUS1*-low TNBC models particularly rely on the MYC transcriptional program. Targeting MYC’s transcriptional activity can be achieved by inhibiting its interaction with its co-activator MAX. Several biomolecule-based or transcriptional MYC-targeting strategies (OMO-103 and OTX-2002) and small-molecule inhibitors of the MYC/MAX interaction (MYCi-361, MYCi-975, and MYCMI-6) have been reported [57–59], providing experimental entry points for testing this therapeutic direction. These agents were selected as representative examples of recent or clinically emerging MYC-targeting strategies, rather than as an exhaustive list of available MYC inhibitors. Among them, OMO-103 is of particular interest since it has already entered phase I clinical testing in advanced solid tumours [57].

#### Targeting DNA_REPAIR to increase replication stress

The `DNA_REPAIR` pathway was up-regulated in *MTUS1*-low TNBC tumours, and contained *MTUS1*-associated essential genes in TNBC cell-line models as well as in the `G2M_CHECKPOINT` and `E2F_TARGETS` pathways. Together, these findings suggest that *MTUS1*-low cells may operate near a replication-stress threshold, where impairment of genome maintenance could promote DNA damage accumulation, premature mitotic entry, and cell mitotic catastrophe (i.e. cell death during mitosis) [66, 67]. The function of *POLR1C* is consistent with this rationale, as it connects RNA polymerase I/III transcriptional stress to genome-maintenance pressure [68]. CX-5461 provides a clinically developed probe of rDNA transcription stress, although it is now also interpreted as a G-quadruplex-stabilising agent whose activity is linked to replication stress and DNA-damage responses [60, 69, 70]. ML-60218 provides a pharmacological tool to perturb RNA polymerase III transcription [61].

#### Targeting UPR to increase ER stress

The `UNFOLDED_PROTEIN_RESPONSE` was enriched both in *MTUS1*-low tumours and among *MTUS1*-associated essential genes, suggesting that proteostasis buffering is critical in *MTUS1*-low TNBC. This is biologically coherent with upregulation of MYC pathways, which increases protein-synthesis load and creates dependence on ER protein folding, glycosylation, and degradation pathways [71]. We selected two candidate targets:*DKC1* and *HYOU1*. For *DKC1*, the pharmacological tool pyrazofurin has been used to target DKC1-linked biology [62], first-in-class HYOU1 inhibitors have been reported [63], which provides pharmacological tools for experimental testing.

## 4 Discussion

One of the most robust results in our study is that *MTUS1*-low TNBC tumours show reproducible enrichment of MYC-linked proliferative pathways compared with their *MTUS1* - high counterparts, although changes in *MYC* expression were modest in most cohorts. This distinction is important because MYC activity can be regulated through transcriptional, post-transcriptional, and post-translational mechanisms [72, 73]. Thus, the data support increased MYC transcriptional activity in *MTUS1*-low TNBC, rather than a model based solely on increased *MYC* mRNA abundance. We also show that this increase is accompanied by up-regulation of the oxidative phosphorylation, and of the unfolded protein response and DNA repair pathways (Figure 2).

Based on these pathways, CRISPR data on TNBC cell line models suggested 3 therapeutic strategies: targeting MYC to reduce its transcriptional activity, targeting the UPR to increase ER stress to untolerable levels, and targeting DNA repair to increase replication stress to untolerable levels. Furthermore, based on CRISPR data, we identified 4 druggable *MTUS1*-associated essential genes (MYC, POLR1C, DKC1 and HYOU1). The corresponding target-inhibitor pairs provided in Table 4 are not presented here as validated treatments for *MTUS1*-low TNBC, but as experimental tools to further evaluate these strategies.

It is important to note that the identification of candidate targets in the present study is limited by the reference pathway database used (Hallmark). Indeed, not all genes are present in the Hallmark, and the exact definition of gene sets involved in a given biological process is somewhat arbitrary. Consequently, interesting candidate targets may have been missed among the list of 87 essential genes. Therefore, we further analysed the functions of these 87 genes, and found three druggable proteins that are absent in Hallmark but whose functions are closely related to UPR: PSMB5, PSMB6 and DPAGT1. DPAGT1 catalyses the first committed step of protein N-glycosylation in the endoplasmic reticulum [74], an important process for maintaining ER proteostasis under folding stress, whereas PSMB5 and PSMB6 are proteasomal subunits that ultimately degrade proteins that failed to fold in the endoplasmic reticulum. Thus, inhibition of PSMB5, PSMB6 or DPAGT1 may lead to cell death based on the same mechanism as DKC1 and HYOU1, i.e. due to an untolerable increase in ER stress, which further supports the interest of this therapeutic strategy for *MTUS1*-low TNBC tumours.

Supplementary Figure places these targets within ER protein processing and illustrates how perturbing distinct steps of this pathway could promote proteotoxic stress. For *DKC1, HYOU1*, and *DPAGT1*, available compounds mainly correspond to preclinical or tool compounds: pyrazofurin [62], HYOU1-directed inhibitors [63], and tunicamycin- or muraymycin-related compounds support DPAGT1 perturbation as a way to interfere with *N*-glycosylation and ER protein processing [65]. By contrast, the proteasome branch is pharmacologically more mature, because clinically approved proteasome inhibitors such as bortezomib and carfilzomib are already available and directly target proteasome activity [75, 76].

*PRMT1* is another example from the list of 87 *MTUS1*-associated essential genes that is not present in Hallmark. It encodes a chromatin- and DNA-damage-associated regulator[77], and CTS2190 is a PRMT1 inhibitor currently being evaluated in an ongoing phase I/II trial in advanced or metastatic solid tumours [64]. Inhibition of PRMT1 may lead to cell death by the same mechanism as POLR1C, i.e. due to an untolerable increase in replication stress. These 4 additional candidate targets (PSMB5, PSMB6, DPAGT1, and PRMT1) were added to Table 4, to illustrate that other candidate targets may also be of interest for the three proposed therapeutic strategies for *MTUS1*-low TNBC. As an example, prior work in our group established that TNBC cell lines in which the *MTUS1* gene has been silenced showed significantly higher sensitivity to WEE1 inhibition and to dual WEE1/PKMYT1 inhibition than their *MTUS1* expressing counterparts [78]. Interestingly, although not present in Hallmark gene sets, the WEE1 and PKMYT1 kinases play a key role at the G2/M checkpoint of the cell-cycle: by inhibiting CDK1, they help to prevent cells from entering mitosis until DNA replication and repair processes are completed [79, 80]. This prior work showed that inhibition of WEE1 or of WEE1/PKMYT1 promotes aberrant mitoses characterised by detachment of centromere proteins from DNA and chromosome pulverisation, leading to cell death during mitosis or after mitotic exit [78].

The examples of WEE1 and PKMYT1 somewhat broaden the rationale of the “Increase in replication stress” therapeutic strategy to “Increase perturbations at the interface between transcriptional stress and genome maintenance”.

Finally, this study presents other limitations related to the datasets used. First, the tumour analyses are retrospective and rely on public cohorts with heterogeneous processing histories, clinical annotations, and sample sizes. Within-cohort stratification and reproducibility criteria across independent datasets were used to reduce this risk, but residual confounding cannot be excluded. Second, the DepMap analysis used endogenous *MTUS1* expression in a limited panel of TNBC cell lines, which are imperfect models of tumours. Selection of candidate targets should therefore be viewed as a prioritisation step to test therapeutic strategies, not as causal evidence that the agressivity of *MTUS1*-low TNBC tumours depends on these individual targets.

## 5 Conclusions

The present study robustly demonstrates that *MTUS1*-low TNBC does not simply mark a subgroup with low expression in *MTUS1*, but it is a biological marker of tumours that share similar biological features: MYC-driven proliferative activity, increased oxidative phosphorylation consistent with higher energy demands, and up-regulation of unfolded protein response and DNA-repair stress-adapation pathways. The main contribution of this work is therefore the definition of a defensible *MTUS1*-low TNBC biological profile. These mechanisms are common to seven independent TNBC cohorts, and are also observed in broader breast cancer cohorts and in several additional TCGA tumour types, which strengthens the proposed profile and argues that it is not a single-dataset artefact. CRISPR data in TNBC cell lines confirmed that *MTUS1*-associated essential genes play functional roles in these pathways, which allowed us to propose 3 therapeutic strategies and their respective candidate target/drug pairs for future experimental validation tests. Of note, prior work in our group on inhibition of WEE1 and PKMYT1 is related to the “Increase replication stress” strategy, which represents a preliminary validation that reinforces the confidence and interest in this strategy.

## Abbreviations

ATIP3: angiotensin II receptor-interacting protein 3
BC: breast cancer
CRISPR: clustered regularly interspaced short palindromic repeats
DepMap: Cancer Dependency Map
ER: endoplasmic reticulum
FDR: false discovery rate
GDC: Genomic Data Commons
GEO: Gene Expression Omnibus
GSEA: gene set enrichment analysis
HER2: human epidermal growth factor receptor 2
MSigDB: Molecular Signatures Database
NES: normalised enrichment score
ORA: over-representation analysis
PR: progesterone receptor
TCGA: The Cancer Genome Atlas
TNBC: triple-negative breast cancer
UPR: unfolded protein response.

## Declarations

### Ethics approval and consent to participate

This study used publicly available, de-identified transcriptomic, clinical annotation, and functional genomics datasets. No new human participants, human tissue samples, or animal experiments were included in this study. Ethics approval and participant consent were obtained by the original studies or data-generating consortia where applicable. No additional ethics approval was required for the present secondary analysis of public data.

### Consent for publication

Not applicable.

### Availability of data and materials

All datasets analysed in this study are publicly available. GEO datasets were retrieved from the Gene Expression Omnibus under accessions GSE181466, GSE192341, and GSE202203. TCGA data were retrieved from the NCI Genomic Data Commons portal (https://portal.gdc.cancer.gov/). SRA-derived datasets SRP042620 and SRP157974 were accessed through recount3. DepMap transcriptomic and Chronos CRISPR–Cas9 gene effect data were obtained from the DepMap Public 24Q4 release through the DepMap portal [35, 36]. Hallmark gene sets were obtained from MSigDB. Derived result tables supporting the conclusions of this manuscript are provided in the article, supplementary material, and accompanying analysis repository.

### Code availability

The analysis code and derived data tables used to generate the publication figures and supplementary dependency analyses are available in the accompanying analysis repository.

