## Supplementary for "Transcriptomic profile of *MTUS1* -low TNBC reveals candidate therapeutic strategies"

### 6 Supplementary

#### 6.1 *MYC* differential expression statistics.

The table S1 reports the differential expression statistics for *MYC* comparing *MTUS1*-low versus *MTUS1*-high samples across TNBC cohorts, broader BC cohorts, and additional TCGA tumour types, including the $\log_2$ fold change (LFC), Wald statistic, and Benjamini–Hochberg adjusted *p*-value.

| **TNBC cohort** | **LFC** | **stat** | **p_adj** |
|---|---|---|---|
| SRP042620 * | 0.461230 | 1.240788 | 0.597948 |
| VUMC * | 0.690917 | 2.426745 | 0.304629 |
| GSE192341 * | 0.040439 | 0.129071 | 0.962640 |
| GSE181466 * | 0.947717 | 3.376026 | 0.010260 |
| TCGA-BRCA * | 0.676129 | 2.555169 | 0.109714 |
| GSE202203 * | 0.821757 | 4.639249 | 0.000017 |
| SRP157974 * | 0.594166 | 5.458394 | $1.504031\times10^{-7}$ |
| Pooled (all TNBC)* | 0.657059 | 8.399523 | $2.209935\times10^{-16}$ |
| **BC cohort** | **LFC** | **stat** | **p_adj** |
| SRP042620 | 0.417268 | 1.376578 | 0.398002 |
| GSE192341 | 0.460685 | 1.699233 | 0.282677 |
| TCGA-BRCA | 0.632592 | 7.406473 | $7.669154\times10^{-13}$ |
| GSE202203 | 0.249000 | 4.319616 | 0.000024 |
| Pooled (all BC) | 0.421463 | 9.925890 | $7.387165\times10^{-23}$ |
| **TCGA cohort** | **LFC** | **stat** | **p_adj** |
| TCGA-KIRC | -0.313257 | -3.234546 | 0.002479 |
| TCGA-KIRP | -0.462571 | -2.765142 | 0.012822 |
| TCGA-BLCA | 0.594900 | 3.592107 | 0.001063 |
| TCGA-COAD | 0.632456 | 7.601203 | $2.374924\times10^{-13}$ |
| TCGA-READ | 0.488680 | 3.316776 | 0.004894 |
| TCGA-PAAD | 0.214854 | 1.443915 | 0.274573 |
| TCGA-LIHC | -0.179705 | -1.049202 | 0.380113 |
| TCGA-LGG | 0.787075 | 7.663400 | $2.055862\times10^{-13}$ |
| TCGA-HNSC | 0.147526 | 1.686403 | 0.132028 |
| TCGA-LUAD | 0.740832 | 6.497252 | $7.115989\times10^{-10}$ |
| TCGA-LUSC | -0.194964 | -1.967787 | 0.118216 |
| TCGA-PRAD | -0.149302 | -1.553031 | 0.152890 |
| TCGA-OV | 0.161193 | 1.314639 | 0.312633 |
| TCGA-UCEC | 0.673513 | 5.388001 | $3.143610\times10^{-07}$ |

Table S1: Differential expression statistics for *MYC* across TNBC cohorts, broader BC cohorts, and additional TCGA tumour types (*MTUS1*-low vs *MTUS1*-high): $\log_2$ fold-change (LFC), Wald statistic (stat), and adjusted *p*-value (p_adj).

#### 6.2 Supplementary DepMap cell lines

| DepMap ID | CCLE name | COSMIC ID |
|---|---|---|
| ACH-000111 | HCC1143_BREAST | |
| ACH-000148 | HCC38_BREAST | 905956 |
| ACH-000212 | BT20_BREAST | 905951 |
| ACH-000223 | HCC1937_BREAST | 905980 |
| ACH-000258 | HCC1187_BREAST | |
| ACH-000276 | HCC70_BREAST | 905962 |
| ACH-000288 | HCC1806_BREAST | 905976 |
| ACH-000374 | MDAMB231_BREAST | 905960 |
| ACH-000573 | HCC1395_BREAST | 905983 |
| ACH-000621 | HS578T_BREAST | 905952 |
| ACH-000624 | SUM159PT_BREAST | |
| ACH-000668 | MDAMB468_BREAST | 905965 |
| ACH-000699 | HCC1599_BREAST | 905977 |
| ACH-000721 | HCC2157_BREAST | 905981 |
| ACH-000768 | CAL120_BREAST | 905953 |
| ACH-000849 | HDQP1_BREAST | |
| ACH-000856 | CAL148_BREAST | |
| ACH-001388 | HCC1500_BREAST | 905963 |
| ACH-001389 | HCC1569_BREAST | 905964 |
| ACH-001390 | HCC1954_BREAST | 905979 |
| ACH-001391 | HCC1428_BREAST | 905982 |
| ACH-001392 | HCC2218_BREAST | |
| ACH-001394 | HCC202_BREAST | |
| ACH-001396 | HCC1419_BREAST | |

Table S2: Metadata for TNBC cell lines used for gene essentiality analyses.

**Table S3:** *MTUS1*-associated essentiality correlations in TNBC cell-line models. Pearson and Spearman correlation coefficients between Chronos gene dependency scores and *MTUS1* expression are reported with unadjusted $p$-values. Positive correlations indicate stronger dependency in lower-*MTUS1* models because more negative Chronos scores reflect stronger essentiality. Values rounded to two significant digits.

| Gene | Pearson $r$ | $p$ (Pearson) | Spearman $\rho$ | $p$ (Spearman) | Min Chronos | Max Chronos |
|---|---|---|---|---|---|---|
| MYC | 0.66 | 0.00046 | 0.66 | 0.00040 | -2.7 | -0.68 |
| DMAP1 | 0.65 | 0.00057 | 0.60 | 0.00190 | -1.7 | -0.52 |
| NOB1 | 0.61 | 0.00160 | 0.61 | 0.00140 | -1.6 | -0.57 |
| DDX49 | 0.61 | 0.00160 | 0.62 | 0.00120 | -2.4 | -0.72 |
| GGPS1 | 0.60 | 0.00200 | 0.62 | 0.00130 | -2.0 | 0.05 |
| DDX10 | 0.58 | 0.00280 | 0.52 | 0.00950 | -2.5 | -0.46 |
| SAP18 | 0.58 | 0.00300 | 0.62 | 0.00120 | -1.4 | -0.41 |
| NOL9 | 0.58 | 0.00320 | 0.60 | 0.00180 | -1.5 | -0.41 |
| RIOK1 | 0.57 | 0.00330 | 0.63 | 0.00100 | -1.5 | -0.50 |
| WDR18 | 0.57 | 0.00340 | 0.53 | 0.00770 | -2.1 | -0.12 |
| SRFBP1 | 0.57 | 0.00340 | 0.62 | 0.00110 | -1.3 | -0.10 |
| URB2 | 0.55 | 0.00560 | 0.52 | 0.00880 | -1.3 | -0.22 |
| THG1L | 0.54 | 0.00620 | 0.64 | 0.00076 | -1.2 | -0.00072 |
| ZNF574 | 0.53 | 0.00730 | 0.55 | 0.00490 | -1.2 | -0.037 |
| IMP3 | 0.53 | 0.00740 | 0.53 | 0.00790 | -2.0 | -0.71 |
| TSR2 | 0.53 | 0.00740 | 0.54 | 0.00610 | -2.4 | -0.97 |
| NOP9 | 0.53 | 0.00770 | 0.53 | 0.00740 | -1.5 | -0.29 |
| UTP4 | 0.53 | 0.00780 | 0.55 | 0.00520 | -2.1 | -0.40 |
| MLST8 | 0.53 | 0.00790 | 0.47 | 0.02100 | -1.5 | -0.12 |

*continued on next page*

| Gene | Pearson $r$ | $p$ (Pearson) | Spearman $\rho$ | $p$ (Spearman) | Min Chronos | Max Chronos |
|---|---|---|---|---|---|---|
| DYNLL1 | 0.52 | 0.00860 | 0.55 | 0.00530 | -1.4 | 0.078 |
| FARSA | 0.52 | 0.00920 | 0.60 | 0.00180 | -1.8 | -0.45 |
| WDR46 | 0.52 | 0.00930 | 0.57 | 0.00380 | -1.8 | -0.39 |
| POP1 | 0.52 | 0.00930 | 0.49 | 0.01600 | -1.4 | -0.77 |
| PPAN | 0.52 | 0.00980 | 0.54 | 0.00700 | -1.6 | -0.67 |
| ABT1 | 0.51 | 0.01000 | 0.58 | 0.00310 | -1.8 | -0.41 |
| ALG1 | 0.51 | 0.01000 | 0.47 | 0.02000 | -1.5 | -0.033 |
| BAP1 | 0.51 | 0.01100 | 0.49 | 0.01600 | -1.0 | 0.0047 |
| RPL14 | 0.51 | 0.01100 | 0.59 | 0.00240 | -2.6 | -1.0 |
| RPL32 | 0.51 | 0.01100 | 0.56 | 0.00410 | -2.9 | -1.6 |
| PSMB6 | 0.51 | 0.01200 | 0.47 | 0.01900 | -2.3 | -0.52 |
| VARS1 | 0.51 | 0.01200 | 0.54 | 0.00640 | -2.3 | -0.86 |
| MSTO1 | 0.50 | 0.01200 | 0.51 | 0.01000 | -2.0 | -0.26 |
| SSB | 0.50 | 0.01300 | 0.56 | 0.00410 | -1.0 | -0.34 |
| FARSB | 0.50 | 0.01300 | 0.55 | 0.00560 | -2.4 | -0.75 |
| GAPDH | 0.50 | 0.01400 | 0.44 | 0.03300 | -1.9 | -0.25 |
| RPL12 | 0.50 | 0.01400 | 0.55 | 0.00520 | -3.3 | -1.5 |
| KRR1 | 0.49 | 0.01400 | 0.55 | 0.00580 | -1.9 | -0.26 |
| RPS16 | 0.49 | 0.01600 | 0.50 | 0.01200 | -2.9 | -1.2 |
| PSMB5 | 0.49 | 0.01600 | 0.46 | 0.02300 | -2.9 | -0.31 |
| OSGEP | 0.48 | 0.01700 | 0.45 | 0.02800 | -1.7 | -0.41 |
| DAD1 | 0.48 | 0.01700 | 0.39 | 0.05800 | -2.1 | -0.66 |
| RSL24D1 | 0.48 | 0.02000 | 0.48 | 0.02000 | -2.6 | -1.2 |
| PWP2 | 0.48 | 0.01800 | 0.51 | 0.01200 | -1.8 | -0.39 |
| EP400 | 0.48 | 0.01900 | 0.44 | 0.03100 | -1.3 | -0.45 |
| AAMP | 0.47 | 0.02000 | 0.51 | 0.01200 | -1.5 | -0.058 |
| SPCS2 | 0.47 | 0.02100 | 0.52 | 0.00990 | -1.2 | -0.51 |
| MASTL | 0.47 | 0.02100 | 0.42 | 0.03900 | -2.1 | -0.20 |
| HIRA | 0.47 | 0.02100 | 0.39 | 0.05900 | -1.3 | 0.090 |
| SLC35B1 | 0.46 | 0.02200 | 0.42 | 0.04300 | -1.8 | -0.24 |
| DHPS | 0.46 | 0.02300 | 0.47 | 0.02000 | -1.6 | -0.32 |
| COPE | 0.46 | 0.02300 | 0.45 | 0.02700 | -1.7 | 0.061 |
| SERBP1 | 0.46 | 0.02300 | 0.38 | 0.06400 | -1.4 | -0.16 |
| RPS15 | 0.46 | 0.02300 | 0.45 | 0.02700 | -2.8 | -1.4 |
| POP7 | 0.46 | 0.02400 | 0.58 | 0.00310 | -1.3 | -0.095 |
| DPF2 | 0.45 | 0.02600 | 0.21 | 0.33000 | -1.6 | -0.25 |
| ARL2 | 0.45 | 0.02700 | 0.52 | 0.00980 | -2.0 | -0.90 |
| ISG20L2 | 0.45 | 0.02700 | 0.42 | 0.03900 | -1.2 | -0.36 |
| PWP1 | 0.45 | 0.02700 | 0.46 | 0.02400 | -1.1 | -0.20 |
| RRP12 | 0.45 | 0.02900 | 0.44 | 0.03000 | -1.8 | -0.74 |
| HYOU1 | 0.44 | 0.03100 | 0.48 | 0.01700 | -1.5 | -0.047 |
| NOL10 | 0.44 | 0.03100 | 0.35 | 0.08900 | -1.8 | -0.52 |
| EIF3I | 0.44 | 0.03100 | 0.44 | 0.03100 | -2.0 | -0.88 |
| RPL37 | 0.44 | 0.03200 | 0.46 | 0.02300 | -2.2 | -0.43 |
| TSR1 | 0.44 | 0.03300 | 0.43 | 0.03400 | -2.0 | -0.64 |
| CHAF1B | 0.43 | 0.03500 | 0.42 | 0.03900 | -2.4 | -1.0 |
| SCD | 0.43 | 0.03500 | 0.34 | 0.09900 | -1.6 | 0.18 |
| PRMT1 | 0.43 | 0.03500 | 0.49 | 0.01600 | -2.2 | -0.53 |
| TTC4 | 0.43 | 0.03700 | 0.34 | 0.10000 | -1.1 | -0.055 |
| POLR1C | 0.43 | 0.03700 | 0.50 | 0.01200 | -2.3 | -0.64 |
| WDR1 | 0.43 | 0.03800 | 0.50 | 0.01300 | -1.7 | 0.074 |
| EIF3F | 0.43 | 0.03800 | 0.41 | 0.04900 | -1.8 | -0.66 |
| MCM6 | 0.43 | 0.03800 | 0.56 | 0.00420 | -1.5 | -0.43 |
| WDR3 | 0.42 | 0.03900 | 0.42 | 0.04000 | -1.7 | -0.63 |
| DPAGT1 | 0.42 | 0.04000 | 0.39 | 0.06000 | -2.3 | 0.20 |
| CHMP4B | 0.42 | 0.04000 | 0.39 | 0.06200 | -2.3 | 0.48 |
| DKC1 | 0.42 | 0.04100 | 0.43 | 0.03600 | -1.8 | -0.63 |
| KAT5 | 0.42 | 0.04200 | 0.39 | 0.05900 | -1.1 | -0.20 |
| EIF2B5 | 0.42 | 0.04300 | 0.44 | 0.03100 | -1.6 | -0.60 |
| CCNK | 0.42 | 0.04300 | 0.41 | 0.04900 | -1.8 | -1.2 |
| NOPCHAP1 | 0.41 | 0.04400 | 0.36 | 0.08600 | -1.0 | 0.088 |
| WDR75 | 0.41 | 0.04500 | 0.44 | 0.03200 | -2.1 | -0.74 |
| POP5 | 0.41 | 0.04500 | 0.42 | 0.04200 | -2.1 | -0.50 |
| TBL3 | 0.41 | 0.04500 | 0.46 | 0.02400 | -1.8 | -0.51 |
| RAB18 | 0.41 | 0.04600 | 0.31 | 0.14000 | -1.1 | 0.57 |
| NOL11 | 0.41 | 0.04900 | 0.40 | 0.05200 | -1.1 | -0.31 |
| ILF2 | 0.41 | 0.04900 | 0.41 | 0.04700 | -1.4 | -0.30 |
| ILF3 | 0.41 | 0.05000 | 0.32 | 0.13000 | -1.7 | -0.39 |

##### 6.3 Protein processing in endoplasmic reticulum.

Because the unfolded protein response was enriched both in *MTUS1*-low tumours and among *MTUS1*-associated essential genes, we designed a simplified version of the protein-processing-in-ER map available in the KEGG database (`https://www.kegg.jp/pathway/map04141`). This figure summarises sequential steps allowing processing of proteins from their entry in the ER, to *N*-glycosylation, chaperone-mediated folding, misfolded-protein handling, and final proteasomal degradation. The *DPAGT1* in *N*-glycan biosynthesis, *HYOU1* in chaperone-mediated folding, and *PSMB5*/*PSMB6* in proteasomal degradation were mapped on this Figure, showing that they play a role at different steps of the response to protein folding.

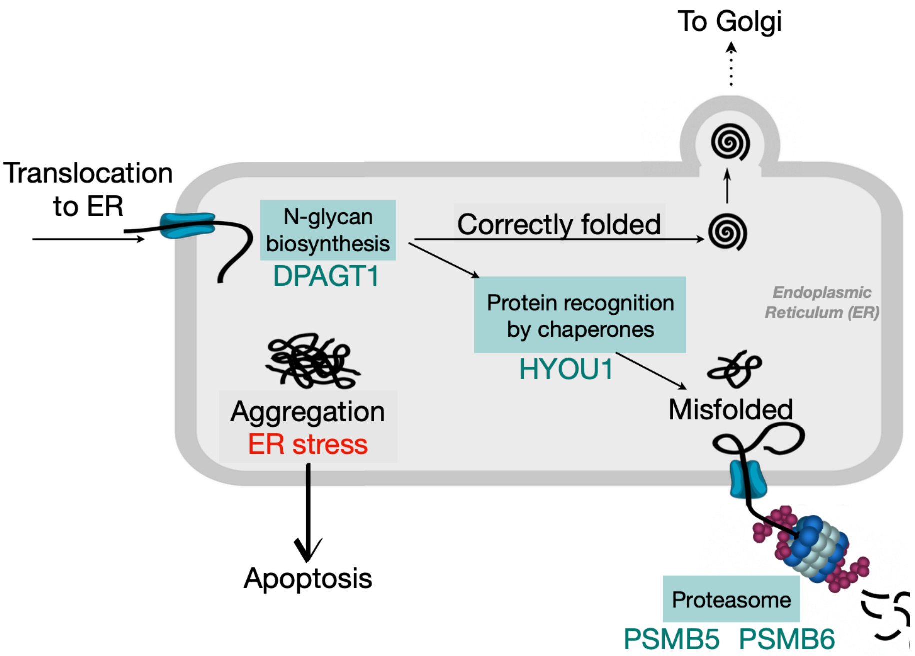


Figure S1: Adapted from the KEGG map *Protein processing in endoplasmic reticulum.* Blue boxes indicate the functions of the four proposed candidate targets whose functions are related to unfolded protein response (DPAGT1, HYOU1, PSMB5, PSMB6).
